# Timing specific parental effects of ocean warming in a coral reef fish

**DOI:** 10.1101/2023.09.27.559693

**Authors:** L.C. Bonzi, R.K. Spinks, J.M. Donelson, P.L. Munday, T. Ravasi, C. Schunter

## Abstract

Population and species persistence in a rapidly warming world will be determined by an organisms’ ability to acclimate to warmer conditions, especially across generations. There is potential for transgenerational acclimation, but the importance of ontogenetic timing in the transmission of environmentally induced parental effects remains mostly unknown. We aimed to disentangle the contributions of two critical ontogenetic stages (juvenile development and adult reproduction) to transgenerational plasticity, by exposing the coral reef fish *Acanthochromis polyacanthus* to simulated ocean warming with natural diel thermal fluctuations across two generations. By using hepatic transcriptomics, we discovered that the developmental environment of the offspring themselves had little effect on their acclimation potential at 2.5 months of life. Instead, the developmental experience of parents increased regulatory RNA production and protein synthesis, which could improve the offspring’s response to warming. Conversely, reproduction in warmer water elicited stress response mechanisms, with suppression of translation and mitochondrial respiration. Mismatches between temperatures in the parental ontogenetic thermal experience deeply affected offspring gene expression profiles, and detrimental effects were also evident when warming occurred both during parents’ development and reproduction. This study reveals that the previous generation’s developmental temperature contributes substantially to thermal acclimation potential during early life, however prolonged heat stress will likely have adverse effects on the species’ persistence.

## Introduction

As a result of anthropogenic climate change, a rise in mean ocean temperatures is happening at an unprecedented rate, and extreme thermal events, such as marine heatwaves, are occurring with increasing frequency and intensity (IPCC, 2023; Oliver et al., 2018). Ocean warming is predicted to impact the distribution and abundance of marine ectothermic organisms (Doney et al., 2012; Hoegh-Guldberg & Bruno, 2010; Scheffers et al., 2016), and tropical species such as coral reef fish might be especially vulnerable to prolonged thermal stress, having evolved in relatively thermal stable environments (Deutsch et al., 2008), as well as living close to their thermal optimum (Rummer et al., 2014). Accordingly, increases in water temperature have shown to cause a rise in oxygen demand (Pörtner, Bock, & Mark, 2017; Schulte, 2015), which, if not met, adversely impacts several traits of coral reef fishes, such as respiratory scope (Nilsson, Crawley, Lunde, & Munday, 2009), swimming ability (Johansen & Jones, 2011), and reproductive output (Donelson, Munday, McCormick, Pankhurst, & Pankhurst, 2010; Spinks, Bonzi, Ravasi, Munday, & Donelson, 2021). However, several fish species have shown the potential to compensate some detrimental effects of ocean warming through acclimation via phenotypic plasticity (Bernal et al., 2018; Bernal, Ravasi, Rodgers, Munday, & Donelson, 2021; Donelson, Munday, McCormick, & Nilsson, 2011; Donelson, Munday, McCormick, & Pitcher, 2012; Grenchik, Donelson, & Munday, 2013; Lee et al., 2020; Salinas & Munch, 2012; Shama, Strobel, Mark, & Wegner, 2014). Switches in energy production mechanisms and substrates, modifications in the protein synthesis machinery, as well as changes in immune and stress responsive gene expression were identified among the major molecular processes underlying such plastic accommodations allowing acclimation to increases in water temperature (Bernal et al., 2018; Bernal et al., 2021; Taewoo Ryu, Veilleux, Donelson, Munday, & Ravasi, 2018; T. Ryu et al., 2020; Shama et al., 2016; Veilleux et al., 2015). Ultimately, thermal plasticity may enable populations and species to persist in a rapidly warming world and offer a lifeline for genetic adaptation to catch up over the longer term.

Acclimation to environmental changes via phenotypic plasticity, however, can be restricted to specific ontogenetic windows of increased sensitivity to external conditions (Burggren & Mueller, 2015). Coral reef fish, as well as other stenothermal species, for example can lack the plasticity to acclimate to increases in temperature during adulthood (Donelson et al., 2010; Nilsson, Östlund-Nilsson, & Munday, 2010), while showing acclimation potential when warming is experienced at development (Donelson et al., 2011; Grenchik et al., 2013). Early life stages are indeed the most sensitive periods to environmental cues and show the highest potential for within-generation plasticity (WGP) (Burton & Metcalfe, 2014; Fawcett & Frankenhuis, 2015; West-Eberhard, 2003), occurring when the organism phenotype is affected by its own experience of the environment. Interestingly, the environmental stimuli perceived during development have been found to not only shape the individual’s phenotype, but also affect subsequent generations via transgenerational plasticity (TGP) (Burton & Metcalfe, 2014). Here, we consider TGP in the inclusive sense of when the experience of previous generations (e.g. parents) influences the offspring phenotype (TGP *sensu* Bell & Hellmann, 2019), and not just when the parental environment interacts with the offspring’s one (anticipatory TGP or TGP *sensu* Salinas, Brown, Mangel, & Munch, 2013). The parental exposure window that is generally thought to have the strongest effect on transgenerational plasticity is during reproduction, because of the temporal proximity and therefore higher cue reliability between the environmental conditions experienced by parents and the future offspring environment (Bell & Hellmann, 2019; Donelan et al., 2020). However, spawning and embryogenesis are also the most vulnerable life stages to high temperatures in fish (Alix, Kjesbu, & Anderson, 2020; Dahlke, Wohlrab, Butzin, & Portner, 2020). Therefore, heat stress during reproduction, for example because of a heatwave, might adversely affect both parents and their offspring, instead of providing the opportunity for beneficial parental effects.

While adaptive parental effects are at the basis of TGP, parental effects that are detrimental to the offspring can also occur. Maladaptive anticipatory TGP, for example, might arise when the environments of parents and offspring do not match, while negative parental carry-over effects can either result from trade-offs between the costs for survival and growth of the parents and the energy invested in reproduction and offspring fitness, or from simple transmission of poor parental condition (Bonduriansky & Crean, 2018; Marshall & Uller, 2007). Californian mussels (*Mytilus californianus*) offspring of thermally exposed parents show reduced tolerance to warming (Waite & Sorte, 2022), and coral reef sea urchin (*Echinometra* sp. *A*) parental exposure to future climate condition negatively affects offspring survival (Karelitz, Lamare, Patel, Gemmell, & Uthicke, 2019). Therefore, the exposure to environmental change across generations does not necessarily lead to acclimation, and the possibility of decreased offspring fitness and detrimental carry-over effects also exist.

The exposure to environmental changes, such as warming, at different ontogenetic timings, and/or in different generations, for example because of heatwaves, could have neutral, additive or interactive effects (Auge, Leverett, Edwards, & Donohue, 2017; Dury & Wade, 2020; Kuijper & Hoyle, 2015; Leimar & McNamara, 2015). In the simplest scenario, a cue elicits the same response regardless of when and how many times it is perceived. In the three-spined stickleback *Gasterosteus aculeatus*, predator-induced WGP and TGP elicit similar phenotypical and molecular responses, with the same set of genes differentially expressed no matter which generation is exposed to the predator cue (Stein, Bukhari, & Bell, 2018). Alternatively, the same cue experienced multiple times could reinforce the information and ensure detection and therefore response (overcoming environmental noise and/or reaching a discrimination threshold) (Bell & Hellmann, 2019), or elicit a stronger, additive response compared to the single experience. In the snail *Nucella lapillus*, when both parents and embryos are exposed to predation risk, offspring size is larger at emergence (Donelan & Trussell, 2018), while additive WGP and TGP effects on growth rate are found in differentially thermally exposed sheepshead minnows (Chang, Lee, & Munch, 2021). Contrasting cues could also be experienced at different times, raising questions on how different experiences interact and are integrated by the organism. For example, the relative importance of the two most sensitive ontogenetic windows in transgenerational plasticity is still debated, and while reproduction at elevated temperatures is sometimes enough to elicit multigenerational thermal TGP (e.g. Lee et al., 2020), in other cases early-life exposure is needed to induce positive parental effects (Radersma, Hegg, Noble, & Uller, 2018). Finally, not all traits have the same plasticity potential, and individual traits might respond differently to the same environmental cue perceived at the same or at different exposure timings (Donelson, Salinas, Munday, & Shama, 2018; Le Roy, Loughland, & Seebacher, 2017; Uller, Nakagawa, & English, 2013). In *Strongylocentrotus purpuratus* sea urchin, DNA methylation shows TGP in response to simulated upwelling conditions, while spicule length only responds to WGP (Strader et al., 2020). Ultimately, the resulting phenotype and acclimation potential of organisms and populations to environmental perturbations will depend on how different traits will be affected and how the environmental cues experienced by parents during the most sensitive windows, development and reproduction, will interact and be integrated with the offspring’s own environmental perception.

Although highly sensitive to warming, the coral reef fish *Acanthochromis polyacanthus* (Bleeker 1855), or spiny chromis damselfish, is able to partially acclimate to warmer temperatures when exposed early in life (developmentally), and to fully restore deficits in aerobic scope caused by higher temperatures when both parents and offspring are exposed to the same warmer conditions (transgenerationally) (Bernal et al., 2018; Bernal et al., 2021; Donelson et al., 2012). A shared suite of differentially expressed genes in the liver, mainly related to lipid, protein and carbohydrate metabolism, immune system and transcriptional regulation, has been found to be related to WGP and TGP mechanisms in this species (Veilleux et al., 2015). Previous studies, however, have always used constant thermal treatments, with no daily fluctuations in temperature mimicking natural variations. Since diurnal variability of environmental stimuli can affect physiological and molecular organism responses (Brakefield & Mazzotta, 1995; Enders & Boisclair, 2016; Kroeker et al., 2020; Schaefer & Ryan, 2006; Schunter, Jarrold, Munday, & Ravasi, 2021), a more accurate imitation of natural conditions could improve predictions of how these fish will respond to global warming. Moreover, until now, experiments on *A. polyacanthus* have continuously exposed parents to warming from hatching to breeding, therefore preventing insights to critical thermal windows for parental exposure to higher temperature that induce TGP. For example, in order to convey beneficial effects on the offspring, parents may only need to experience warming as adults during gametogenesis and reproduction, or alternatively, they may need to be exposed to warmer conditions during development in early life. While exposure timing within the F2 generation was explored in Bernal et al. (2021), there are some limitations as one of the orthogonal F3 crosses is missing and both the F1-F2 generation were exposed continuously before testing the impacts of reproductive thermal exposure. Therefore, further investigation into the role of warming during parental developmental and reproductive stages is needed to improve our understanding of the interplay between the timing of environmental variation and plasticity. Finally, since reproductive exposure in this species also coincides with embryogenesis because of nest care, the independent exposure of parents to elevated temperature during development or during reproduction only is necessary to determine if the transgenerational acclimation effects reported so far are indeed parental effects or rather offspring developmental plasticity due to embryo exposure to warming.

In this study, we exposed F1 *A. polyacanthus* to daily and seasonally fluctuating present-day control or +1.5°C average increased temperature during development and at reproduction (Fig. 1). F2 offspring clutches from these parents were split post-hatching into control or +1.5°C, where they developed for 80 days. Elevated temperature exposure at reproduction caused low reproductive output and poor hatchling quality (Spinks et al., 2021). At 80 days post-hatching, offspring from parents that reproduced at elevated temperature were still in poorer body condition compared to offspring from control parents, regardless of the parental and of the offspring’s own developmental thermal environment (Spinks, Donelson, Bonzi, Ravasi, & Munday, 2022). Additionally, parental exposure to elevated temperature at development also resulted in lower 80-day old offspring condition. Parental exposure to elevated temperature, irrespective of ontogenetic timing, seems therefore to be causing maladaptive effects in the offspring. Alternatively, reduced weight might represent a trade-off with the adaptive parental effects on metabolism previously recorded for this species (Bernal et al., 2021; Donelson et al., 2012; Munday, Donelson, & Domingos, 2017). Here, we assessed the genome-wide liver gene expression of the F2 offspring raised for 80 days at elevated or control temperature from all the parental combinations. The 184 analysed transcriptomes allowed us to explore the molecular responses of *A. polyacanthus* to warming at different ontogenetic timings over two generations and disentangle the effects of the parental experience during development from the reproductive and the offspring developmental exposures. Because of the concurrent increase in mean ocean temperatures as well as more frequent and intense heatwaves, this study aimed to answer unresolved questions related to transgenerational acclimation potential to ocean warming, especially taking into account the possibility for temperature mismatches between generations and also during an individual’s lifespan.

## Methods

### Experimental design

In order to investigate the importance of exposure timing in the response to warming, we analysed liver gene expression of *A. polyacanthus* exposed to elevated temperature over two generations. Detailed descriptions of the experimental set-up are provided in Spinks et al. (2021; 2022). Briefly, adult spiny chromis damselfish (F0 generation) were collected from the wild in the Palm Islands region (18°37′ S, 146°30′ E) and nearby Bramble Reef (18°22′S, 146°40′E) of the central Great Barrier Reef in Australia, paired and housed with seasonally cycling water temperature resembling the collection site conditions. During the Austral summer of 2016, F0 pairs bred, and egg clutches were kept with parents until hatching, in order for them to provide nest care as in the wild. To account for genotypic variation, newly hatched F1 siblings from six breeding pairs were split between a present-day control and an elevated temperature treatment. Water temperature of the two thermal treatments was finely controlled to match simulated seasonal (winter minimum 23.2°C, summer maximum 28.5°C) and diurnal (3:00 am -0.6°C, 3:00 pm +0.6°C) cycles of the Palm Islands for the control treatment, with the elevated thermal treatment matching this, but with an increase of 1.5°C. This temperature regime was chosen to match the projections for average ocean temperature increase by 2100 (Fox-Kemper et al., 2021), and already occurring heatwaves (Frölicher, Fischer, & Gruber, 2018). F1 fish were kept in the two thermal treatments until maturity (∼1.5 years of age). In the late Austral winter of 2017, fish from different families were paired for reciprocal sex crosses of the developmental temperatures, and the formed pairs were further placed into present-day control or +1.5°C reproductive temperatures (Fig. 1). The F2 generation was produced between December 2017 and April 2018, although no reproduction occurred for the elevated reproductive thermal environment when both parents developed at +1.5°C. Since eggs were kept with their parent until hatching, embryogenesis occurred at reproductive temperature. Newly hatched F2 generation siblings were split into present-day control or +1.5°C thermal treatments (Fig. 1), which followed the above-mentioned seasonal and diurnal cycles of water temperature. During rearing, approximately 9% natural mortality occurred (Spinks et al., 2022). At 80 days post-hatching (dph), F2 fish were sexed by external examination of their urogenital papilla, and two males and two females per clutch per treatment (twelve to twenty individual fish per treatment; Suppl. Table S1) were euthanized by cervical dislocation, measured for standard length, weighed and dissected. Livers were immediately snap-frozen in liquid nitrogen and stored at -80°C for subsequent RNA extraction. Liver was chosen as the target tissue of this study because of its major role in metabolism and to allow comparisons with previous works (Bernal et al., 2018; Bernal et al., 2021; Taewoo Ryu et al., 2018; T. Ryu et al., 2020; Veilleux et al., 2015). All samples were collected between 9 am and 12 pm. All procedures were performed in accordance with relevant guidelines and were conducted under James Cook University’s animal ethics approval A1990, A2210 and A2315.

**Figure 1.**
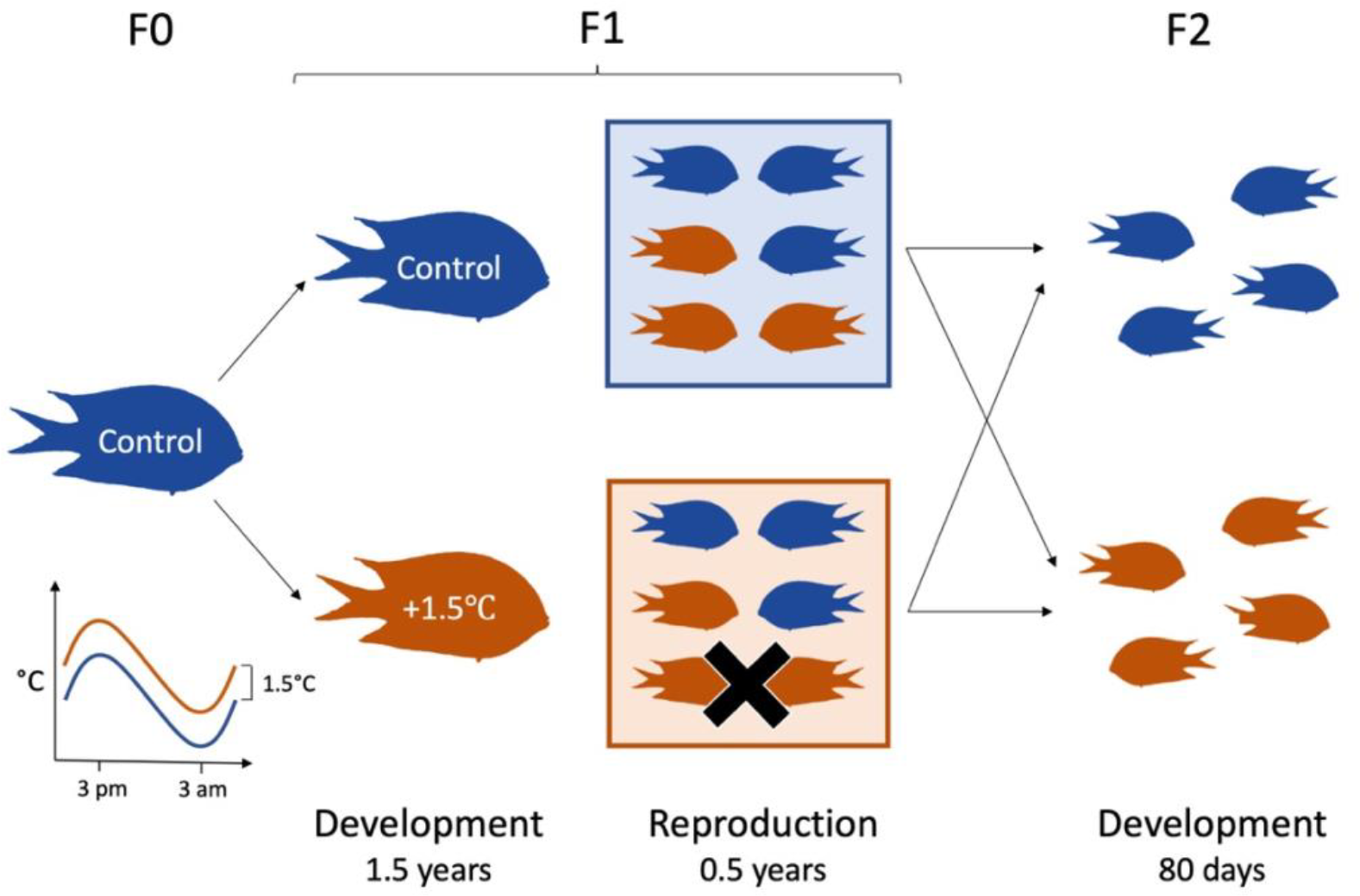
Experimental design. F1 *A. polyacanthus* from wild-caught F0 developed either at present-day control temperature (control - blue) or elevated temperature (+1.5°C - orange) with seasonal and daily fluctuations. At 1.5 years of age, F1 were paired in reciprocal sex crosses of the two thermal treatments and further exposed to control (blue rectangle) or elevated (orange rectangle) reproductive temperatures. The black “X” indicates the F1 treatment that did not reproduce. F2 siblings were split after hatching into control or elevated temperatures, where they developed for 80 days.

### RNA sequencing

Frozen livers were homogenized in Qiagen RLT Plus buffer for 30s with single use silicone beads in an MP Biomedicals FastPrep-24 homogenizer. Total RNA was isolated from whole homogenized livers using a mirVana miRNA Isolation kit, following the manufacturer’s protocol for total RNA isolation procedure. A Nanodrop (Thermo Scientific) and a 2100 Bioanalyzer (Agilent) were used to determine quantity and quality of the isolated RNA. RNA-Seq libraries were prepared using the Illumina TruSeq stranded mRNA Library Preparation Kit, with each sample uniquely barcoded. Library quality check and quantification were performed with a Bioanayzer High Sensitivity DNA assay (Agilent). Paired-end fragments of 150 base pairs were sequenced with an Illumina HiSeq 4000 at the King Abdullah University of Science and Technology Bioscience Core Lab. Samples from different thermal treatments were randomly assigned to each lane to avoid batch effects during sequencing, for a total of 184 sequenced samples (Suppl. Table S1).

### Gene expression analyses

Read quality check was performed with FastQC (Andrews, 2010) before and after quality trimming and adapter removal by Trimmomatic v0.39 (Bolger, Lohse, & Usadel, 2014) with parameters: “TRAILING:3 SLIDINGWINDOW:4:15 MINLEN:40”. Trimmed reads were mapped against the *Acanthochromis polyacanthus* genome (ENSEMBL ASM210954v1) using HiSAT2 v2.2.1 (Kim, Paggi, Park, Bennett, & Salzberg, 2019) with default settings, providing a list of known splice sites and specifying strand-specific information (--rna-strandness = RF). The featureCounts function from the Subread v2.0.2 package (Liao, Smyth, & Shi, 2014) was used to calculate gene counts, in read pair counting mode, allowing for multimapping fractional computation.

The DESeq2 package v1.26.0 (Love, Huber, & Anders, 2014) was used to statistically analyse differential gene expression in R v3.6.3 (R Core Team, 2020). The presence of outliers and batch effects in the data was evaluated through clustering and visualization using variance stabilized transformed (VST) counts. Based on principal component analyses (PCAs) and heatmaps of the sample-to-sample distances, six outlier samples were identified and excluded from further analyses (Suppl. Table S1). Likelihood ratio tests (LRTs) were used to determine the best models. The chosen design formula included the variable “family” to control for differences due to the parental lineage, and the main effects of developmental, reproductive and offspring developmental thermal environments, as well as the interaction between parental developmental and reproductive temperature: ∼Family + Parental pair development * Reproduction + Offspring development, since no significant interaction was found with offspring developmental temperature. Offspring gender was not found to be influential and was therefore excluded from the model. Differentially expressed genes (DEGs) were statistically determined for the main effects and their interactions, using false discovery rate (FDR) adjusted p-value < 0.01 (Benjamini & Hochberg, 1995) and a mean expression of > 10 reads (baseMean) as threshold. LRT identified DEGs due to the interaction found between parental development and reproduction were analysed using the degPatterns function from the DEGreport R package v1.22.0 (Pantano, 2021) to identify clusters of differentially expressed genes with similar expression profiles. The function was run on the VST processed count matrix of such genes with default settings, except for cluster outlier removal (reduce = TRUE) and minimum number of genes per cluster set to 50 (minc = 50). Wald tests were used to run pairwise comparisons between treatments, and DEGs identified by the following cut-offs: adjusted p-value < 0.01, apeglm shrunken |Log2 Fold Change (log2FC)| ≥ 0.3 to reduce false positives (Zhu, Ibrahim, & Love, 2019), and baseMean > 10.

Additionally, R package WGCNA v1.70-3 (Langfelder & Horvath, 2008) was used to run weighted gene correlation network analyses in order to identify clusters (modules) of highly correlated genes and relate them to the experimental treatments and offspring traits. The analysis was run using VST counts of genes with average counts >1 in more than eleven samples, which is the smallest sample set per treatment after outlier removal. A soft-thresholding power β of 7 was chosen based on network topology analysis, and gene network clusters were identified using the automatic one-step network construction and module detection blockwiseModules function, using a signed topological overlap matrix (TOM), a minimum module size of 30 and a threshold of 0.3 for merging modules. Gene modules were then correlated with the parental, reproductive and offspring thermal treatments, sex, standard length and weight, to identify which gene clusters were significantly associated with the different thermal exposure timings (p-value < 0.001). A module-trait relationships heatmap was produced and modules with significant correlations further investigated.

Functional enrichment analyses of differentially expressed genes identified by DESeq2, as well as gene sets belonging to degPatterns clusters and to significant modules detected by WGCNA were performed in OmicsBox v2.0.36 (BioBam Bioinformatics, 2019) with Fisher’s Exact Test (FDR < 0.05).

## Results

Exposure of *A. polyacanthus* to elevated water temperature at different ontogenetic times greatly affected the 80-day old juvenile hepatic gene expression profiles. The strongest driver of gene expression variation in the offspring livers was the thermal environment in which their parents either developed (4,018 differentially expressed genes, DEGs; Fig. 2; Suppl. Table S2) or reproduced (4,065 DEGs; Fig. 2; Suppl. Table S3). Conversely, the offspring post-hatching developmental temperature had the smallest influence on their own expression profile (1,483 DEGs; Fig. 2; Suppl. Table S4). Only a relatively small number of genes (274; Fig. 2; Suppl. Table S5) was influenced by elevated temperature irrespective of the exposure timing (parental development, reproduction and offspring development) and they were mostly involved in metabolism and oxidoreductase activity, including cytochrome P450 (CYP) superfamily of enzymes.

**Figure 2.**
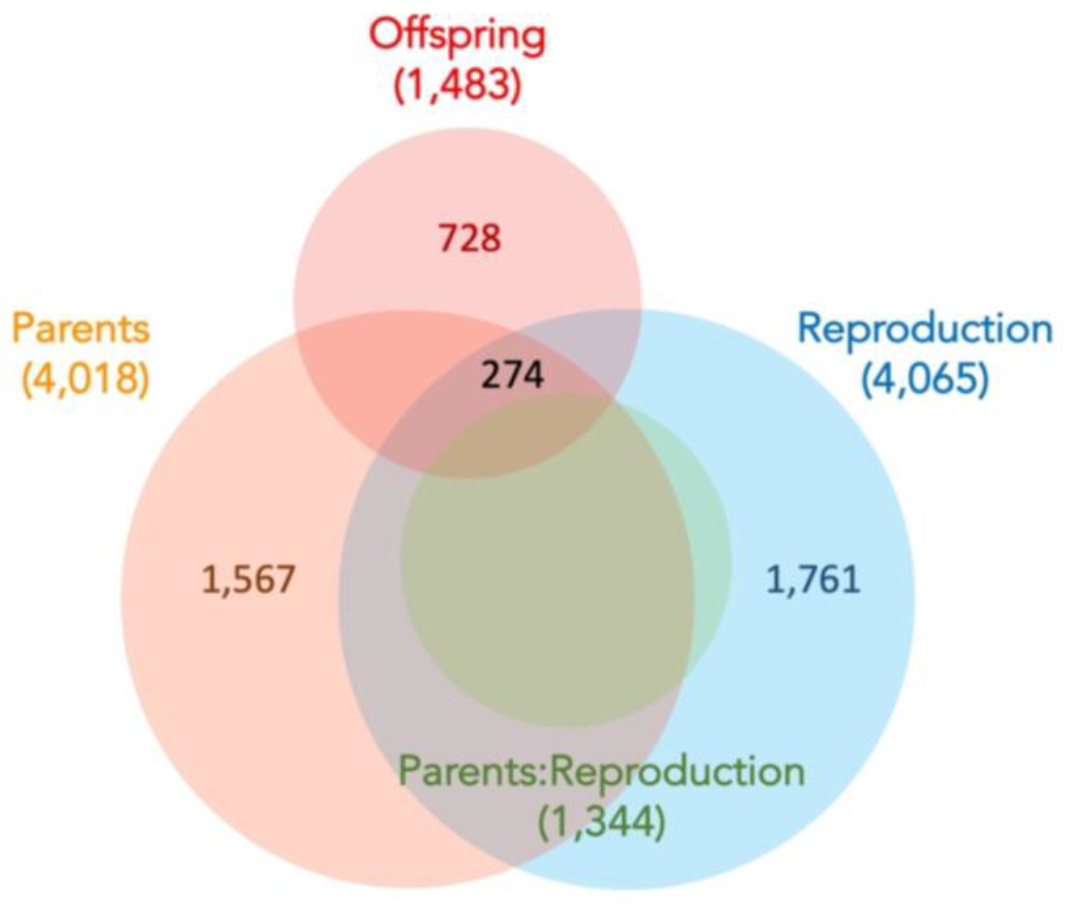
Offspring genes significantly affected by warming at different timing. Venn diagram of hepatic F2 offspring genes whose expression is influenced by the three main effects, parental developmental (Parents), reproductive (Reproduction), and offspring developmental (Offspring) thermal experiences, and by the interaction between the parental developmental and reproductive temperatures (Parents:Reproduction), as identified by likelihood ratio tests.

### Effects of warming during offspring development

The temperature experienced by offspring at development explained changes in expression levels of 728 genes, which were not altered due to any parental exposure timing (Fig. 2; Suppl. Table S4) and were mainly involved in DNA replication and repair and tRNA aminoacylation for protein translation (Suppl. Table S6). When offspring were exposed to elevated temperature, regardless of their parental developmental and reproductive temperatures, genes encoding for components of the MCM complex (MCM2, MCM3, MCM4, MCM5, ZMCM6B) involved in DNA replication initiation were downregulated, as well as CYP coding genes involved in oxidoreductase activity (Suppl. Table S7). Accordingly, genes involved in DNA replication initiation and DNA repair belonged to a gene network cluster significantly negatively correlated with offspring developmental temperature (64 genes; p-value 6e^-06^; Suppl. Fig. S1 & S2; Suppl. Table S8). No interaction between the offspring’s developmental thermal environment and any of the parental thermal exposures was found, therefore offspring response to elevated temperature did not vary significantly if matched or mismatched in temperature with any of the parental thermal environments.

### Transgenerational warming effects

The parental thermal exposure, either during development or reproduction, affected almost four-times the number of genes compared to the offspring’s own developmental thermal environment (Fig. 2). Accordingly, the largest differences in gene expression were found between offspring with different parental developmental and/or reproductive thermal experiences, while comparing between offspring raised at different temperatures usually returned the smallest DEG numbers (Fig. 3; Suppl. Fig. S3).

**Figure 3.**
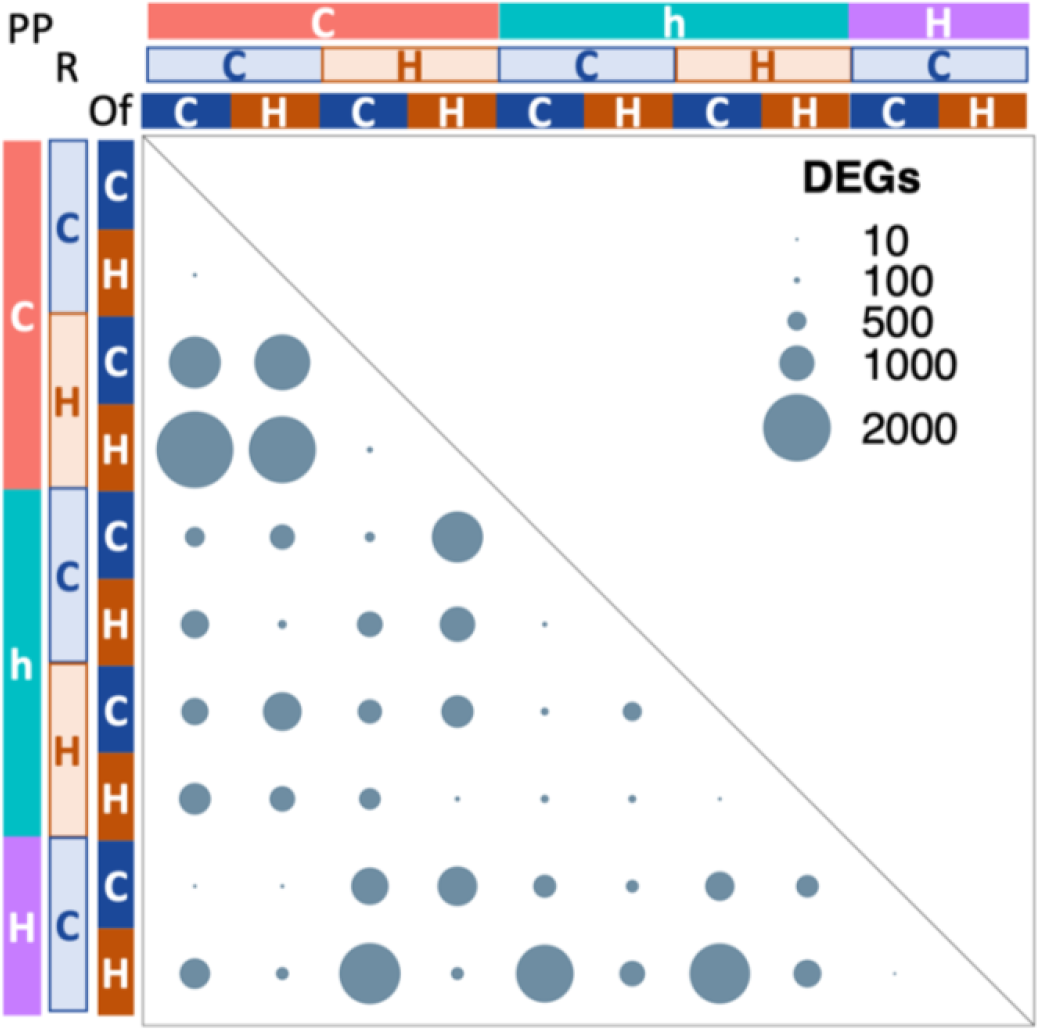
Offspring differentially expressed genes (DEGs) due to different thermal treatments. DEGs from pairwise comparisons between offspring with contrasting thermal histories. “PP” stands for Parental Pair developmental, where “C” = both parents developed at control temperature, “h” = one parent developed at control and one parent developed at +1.5°C, “H” = both parents developed at +1.5°C. “R” stands for Reproductive and “Of” for Offspring developmental thermal conditions, where “C” = control temperature, “H” = +1.5°C. The size of the circles is proportional to the number of DEGs between comparisons (FDR < 0.01).

We found a strong interactive effect between parental developmental and reproductive thermal experiences, with 1,344 genes (Fig. 2; Suppl. Table S9) differing in expression if there was a mismatch between the temperature at which parents developed and the temperature at which they reproduced. Proteolysis, especially proteasome-mediated ubiquitin-dependent protein catabolic process, as well as protein folding, cell redox homeostasis and peptide biosynthesis were among the processes differentially regulated whenever a mismatch of the two parental exposure temperatures occurred (Supp. Table S9). These functions were mostly associated with a cluster of 427 genes that showed increased expression in offspring from parents that experienced temperature mismatch during development and reproduction, regardless of the offspring developmental temperature (Fig. 4A; Suppl. Table S10). Within this cluster, differential expression was also related to DNA double-strand break repair and protein folding quality control in the endoplasmic reticulum. For example, components of the unfolded protein response (UPR) pathway, such as heat shock protein family A member 5 (HSPA5), DNAJ heat shock protein family B11 (DNAJB11) and zinc finger and BTB domain containing 17 (ZBTB17), as well as members of the calnexin/calreticulin cycle, like calnexin (CANX), calreticulin (CALR), protein disulphide isomerase family A members 3 and 4 (PDIA3, PDIA4), UDP-glucose glycoprotein glucosyltransferase 1 and 2 (UGGT1, UGGT2), and protein kinase C substrate 80K-H (PRKCSH), were all found in this cluster of genes. Further processes such as ATPase-coupled transmembrane transporter and phosphopyruvate hydratase activities were conversely downregulated only if parents were exposed to elevated temperature either at development or at reproduction alone (348 genes; Fig. 4B; Suppl. Table S11), while genes involved in circadian rhythm were strongly downregulated in offspring from parents exposed to warming at reproduction alone, but slightly upregulated if elevated temperature was experienced during parental development only (52 genes; Fig. 4C; Supp. Table S12).

**Figure 4.**
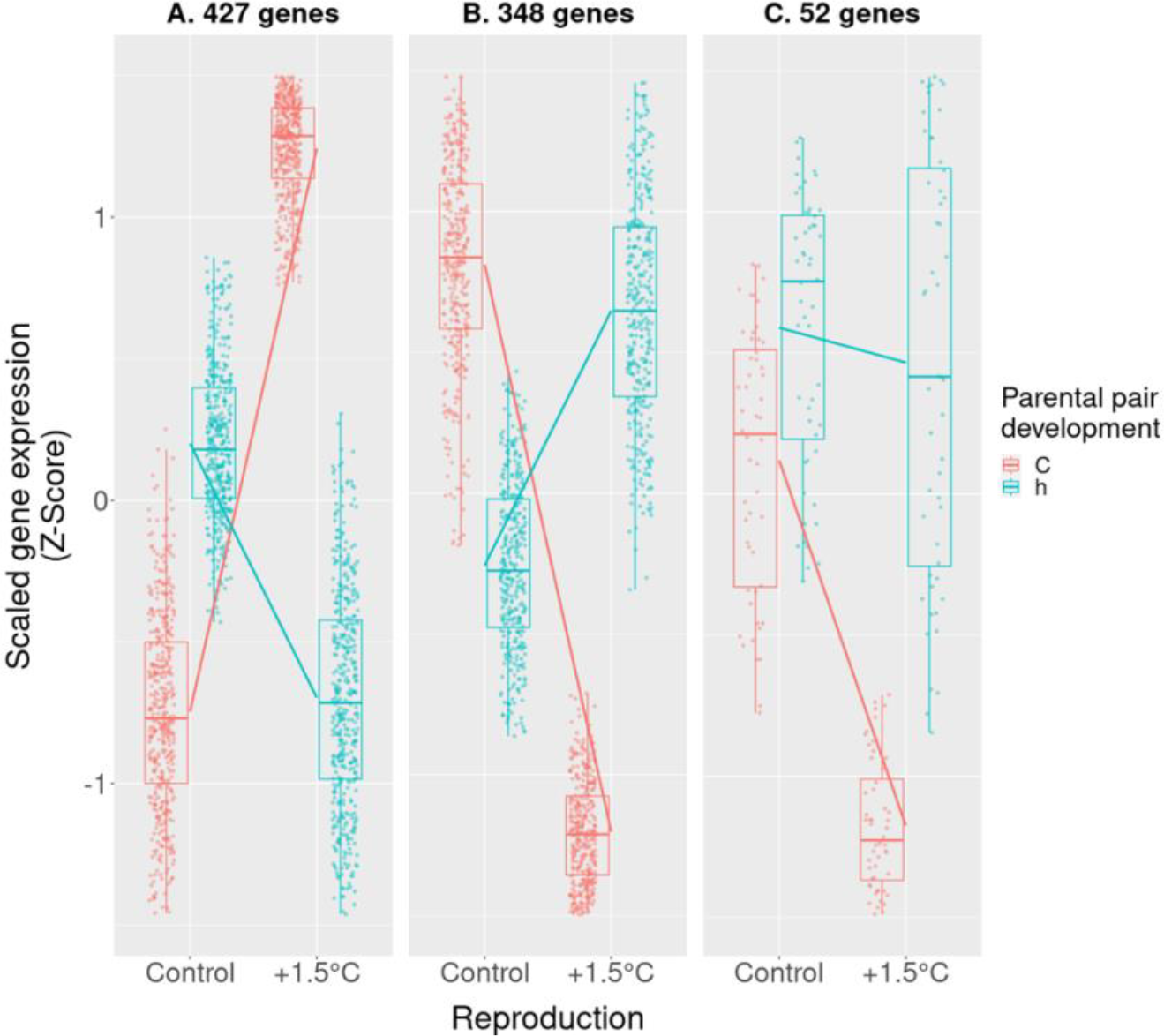
Expression profiles of differentially expressed genes (DEGs) due to the interaction between the two parental exposure timings. DEGs were clustered based on their scaled expression profiles (Z-score). The number of genes per cluster is provided above each plot. In the parental pair development, “C” = both parents developed at control, “h” = one parent developed at control and one at +1.5°C.

### Effects of warming during parental development

The temperature experienced by parents during development, and not at any other exposure timings, caused expression changes in 1,567 offspring genes (Fig. 2; Suppl. Table S2), mostly involved in functions such as DNA-directed 5’-3’ RNA polymerase activity, gene expression, tRNA processing, ribonucleoprotein complex biogenesis and RNA splicing (Suppl. Table S13). Several RNA polymerase I and III subunits (POLR1A, POLR1B, POLR1D, POLR1E, POLR3A, POLR3C, POLR3D) and genes with rRNA and tRNA processing functions in particular were upregulated in offspring of parents that developed at elevated temperature and reproduced at control, regardless of the offspring own developmental temperature (Suppl. Tables S14 & S15). The downregulated genes showed functions related to vitamin B6 binding, including genes involved in glucose/energy metabolism such as glutamic--pyruvic transaminase (GPT), glutamic-oxaloacetic transaminase 1 (GOT1), serine hydroxymethyltransferase 2 (SHMT2), cystathionine gamma-lyase (CTH) and glycogen phosphorylase L (PYGL), as well as steroid hormone mediated signalling, due to several nuclear receptors (NR0B2, NR1D2, NR1I2, NR2F6, NR3C1, RORA, RXRG) (Suppl. Table S14 & S15). If only one of the parents developed at elevated temperature, however, their offspring downregulated genes encoding for structural constituents of ribosomes and involved in translation and peptide biosynthesis, while among the upregulated genes we found genes encoding for endoplasmic reticulum proteins such as CALR and torsin family 1 member A (TOR1A), involved in quality control of protein folding, as well as genes with lipid metabolic functions (Suppl. Tables S16 & S17).

### Effects of warming at reproduction

Translation and cellular respiration were the main processes altered in the 1,761 offspring genes exclusively differentially expressed because of warming at reproduction (Fig. 2; Suppl. Tables S3 & S18). Downregulation due to elevated temperature at reproduction was found for structural constituents of ribosome and genes with peptide synthesis function, as well as oxidoreductase activity, ATP synthesis and electron transfer (e.g. cytochrome-c oxidase and NADH:ubiquinone oxidoreductase subunits), regardless of either parental or offspring developmental temperature (Suppl. Tables S19-22). Accordingly, genes coding for structural constituents of ribosomes involved in translation were found in a gene network cluster significantly negatively correlated with reproductive temperature (p-value 2e^-12^; 812 genes; Suppl. Fig. S4A; Suppl. Table S23), while genes related to electron transfer activity, cellular respiration, translation, proteasome complex, and mRNA splicing were part of a larger cluster of 2,432 genes also negatively correlated with reproductive temperature (p-value 9e^-04^), as well as with offspring length (p-value 4e^-05^) and weight (p-value 2e^-07^) (Suppl. Fig. S4B; Suppl. Table S24). Differences were instead found in the upregulated functions depending on if any of the parents was also exposed to elevated temperature during development. If both parents were exposed to warming at reproduction only, their offspring upregulated genes related to translation initiation and regulation (e.g. translation initiation factor 2 alpha kinases 2 and 3 - EIF2AK2, EIF2AK3 - and 4E binding protein 2 - EIF4EBP2), protein folding (e.g. TOR1A), and negative regulation of gene expression. Moreover, many of these genes encoded for proteins localized in the endoplasmic reticulum, for example genes in the calnexin/calreticulin cycle (CALR, CALX, PDIA3, PDIA4) and in the ubiquitin-dependent ERAD pathway (ER lipid raft associated 2 - ERLIN2), as well as UPR pathway components (signal sequence receptor subunit 1 - SSR1, endoplasmic reticulum oxidoreductase 1 alpha - ERO1A, DNAJB11, PDIA5, PDIA6, ZBTB17) (Suppl. Table S19 & S20). Genes located in the ER and involved in protein folding were also found in a small module positively correlated with reproductive thermal treatment (164 genes; p-value 3e^-07^; Suppl. Fig. S4C; Suppl. Table S25). On the other hand, if one of the parents not only reproduced but also developed at elevated temperature, their offspring overexpressed genes involved in protein transport, regulation of systemic arterial blood pressure by circulatory reninangiotensin (angiotensin converting enzyme 2 - ACE2, glutamyl aminopeptidase - ENPEP), protein ubiquitination and apoptosis (Suppl. Table S21 & 22).

## Discussion

In this study, we show the conserved and timing-specific molecular signatures of exposure to near-future predicted water temperatures across two generations and reveal that the parental thermal experience has a much greater effect on the offspring transcriptional response at 80 days old than does their own posthatching developmental experience. This builds our understanding on the influence of environmental conditions across generations and supports the expectation that parental influence is likely to be strongest in early life (Yin, Zhou, Lin, Li, & Zhang, 2019), while current environmental conditions become the main driver later in development (e.g. at one year in Bernal et al., 2021). Accordingly, we observed in this experiment that offspring developmental temperature had a significant effect on their own body condition only when their parents developed and reproduced at ambient temperature, with the parental thermal exposure otherwise controlling offspring length and weight ratio (Spinks et al., 2022). The consistency between parental and offspring thermal exposures did not affect offspring condition (Spinks et al., 2022) or molecular responses, suggesting carry-over rather than anticipatory parental effects (Bonduriansky & Crean, 2018; Uller et al., 2013). On the contrary, mismatches between the temperatures experienced by the two parents at development or across parental lifespan played an important role in shaping offspring thermal acclimation.

The exposure of parents and offspring to higher temperature caused the activation of some common molecular responses. In particular, we found modifications in metabolism and energy production, a common response to warming in ectotherms such as fishes, also found in other ocean warming experiments, both within (Bernal et al., 2020; Smith, Bernatchez, & Beheregaray, 2013) and across generations (Bernal et al., 2018; Bernal et al., 2021; Veilleux et al., 2015). Members of the cytochrome P450 superfamily of enzymes, characterized by oxidoreductase activity, were downregulated whenever offspring or their parents were exposed to elevated water temperature. Downregulation of hepatic CYP mRNAs, proteins and activity is usually linked to inflammation states in animal models (Aitken, Richardson, & Morgan, 2006; Stavropoulou, Pircalabioru, & Bezirtzoglou, 2018), and hence our results suggest liver inflammation in offspring directly or transgenerationally exposed to warming. While in previous studies inflammatory states have been found in acutely exposed fish only (Bernal et al., 2018; Veilleux et al., 2015), our experimental setup introduced for the first time a diel temperature fluctuation which might aggravate the negative effects of warming. These results are also supported by our measurements of reproductive output (Spinks et al., 2021) and the lack of reproductive success in pairs where both individuals were exposed to warming throughout their life. Similarly, diurnal fluctuations exacerbated the effects of temperature in reducing fathead minnows *Pimephales promelas* size, compared to constant warming, despite having beneficial effects on their upper thermal tolerance limits (Salinas, Irvine, Schertzing, Golden, & Munch, 2019). Notably, in the fruit fly *Drosophila melanogaster*, the same increased thermal tolerance provided by temperature fluctuations was however accompanied by fitness and reproductive costs, such as lower fecundity and reproductive output (Cavieres et al., 2020). Yet, since we lack a non-fluctuating treatment within our experiment, we cannot rule out the possibility that the differences between this and previous studies are due to other factors, and not the diel fluctuation in temperature. Nevertheless, to better evaluate the effects of transgenerational plasticity, and more generally to accurately predict organism responses to climate change, it is crucial to incorporate temporal fluctuations that resemble the natural environment in the best way possible.

The exposure of the two generations to elevated temperature mostly caused the activation of distinct gene sets in the offspring, similarly to the discordant within- and across-generation transcriptional responses of *Daphnia ambigua* in a predator-induced phenotypic plasticity experiment (Hales et al., 2017). Exclusive to the offspring developmental warming experience was the inhibition of DNA replication and the activation of DNA repair mechanisms. Suppression of DNA replication because of developmental warming might be a sign of the activation of the DNA-replication stress-response pathway (Osborn, Elledge, & Zou, 2002) to allow time to repair damaged DNA following temperature stress. Alternatively, DNA replication downregulation could indicate energy investment shifts, in agreement with findings from Bernal et al. (2018), where *A. polyacanthus* offspring downregulated genes involved in DNA replication concurrently with heightened routine oxygen consumptions when developmentally exposed to an increase in water temperature compared to the previous generation. By contrast, parental thermal exposure affected processes related to RNA processing and protein synthesis, modifications and catabolism, indicators of changes in protein turnover, possibly a conserved transgenerational response to warming found in *A. polyacanthus* as well as in sticklebacks (Shama et al., 2016; Veilleux et al., 2015). Hence, a combination of shared and distinct mechanisms underlying inter-generational and post-hatching thermal plasticity are likely in place, revealing a core response to warming but also a decoupling in the two exposure timing effects which could potentially be independently subject to evolutionary pressure (Bell & Stein, 2017).

The two different parental exposure timings, either throughout juvenile development until maturity or during reproduction only, also elicited distinct molecular responses in the offspring. Likewise, in sticklebacks, grandparental reproductive and parental developmental exposures elicit different physiological responses in subsequent generations (Shama et al., 2016; Shama et al., 2014; Shama & Wegner, 2014). Here, the parental developmental exposure to elevated temperature caused changes in energy utilization in offspring through downregulation of genes involved in glucose metabolism and nuclear hormone receptors like NR1D2 and RORA, which are key regulators of the circadian clock and many metabolic functions (Cho et al., 2012; Yang, Lamia, & Evans, 2007). Parental developmental warming also increased transcription and modification potential of non-coding RNAs, in particular rRNAs and tRNAs, which may indicate heightened protein synthesis as well as cell growth (Goodfellow & Zomerdijk, 2013; Turowski & Tollervey, 2016). Interestingly, sticklebacks born from mothers developmentally exposed to warming revealed enhanced protein synthesis resulting in higher respiration rates, and, consequently, in the ability to meet the increased oxygen demand in warmer water and maintain aerobic scope (Shama et al., 2016). Because *A. polyacanthus* has been similarly found to retain aerobic capacity when transgenerationally exposed to warming (Donelson et al., 2012), our results suggest that increased protein synthesis is a conserved molecular mechanism underlying the acclimation ability in this and other species. Moreover, our findings reveal that such beneficial traits are due to the developmental exposure of parents to increased water temperatures, therefore being transgenerational responses rather than WGP due to embryo exposure to warming. The beneficial effects of developmental parental exposure to elevated temperature were also evident in the ability of 80-day old offspring to maintain swimming performance (Spinks, 2021), despite being lighter and in lower body condition compared to offspring from control parents (Spinks et al., 2022). Trade-offs seem therefore to be in place between benefits and costs of the metabolic adjustments need to acclimate to elevated temperature. Nevertheless, parental exposure of *A. polyacanthus* to warming during development appears to be overall beneficial for the offspring and might lead to improved acclimation to increased water temperature.

The benefits of parental development on offspring acclimation potential seemed however to be reduced when only one parent was exposed to elevated temperature during development, while the other developed at control temperature. Indeed, offspring from developmentally mismatched parents exhibited signs of metabolic stress, impairment of the translational machinery, and maladaptive swimming speed (Spinks, 2021). Our findings therefore suggest that the adaptive nature of the transgenerational effects due to the developmental exposure timing might depend on the consistency between parental thermal experiences during early life. Moreover, while here we cannot disentangle the individual effects of each parent, paternal and maternal contributions to the offspring acclimation potential might also differ, and future research focused on the individual parental contributions is needed to fully disentangle each parent’s influence on the new generation’s persistence at elevated temperature.

While parental developmental thermal exposure caused trade-offs between costs and benefits of thermal acclimation in the offspring, the exposure to elevated temperature during reproduction, which in our experiment coincided with both gametogenesis and embryo development, was always detrimental for the offspring, and caused a marked reduction in expression of genes involved in protein synthesis and mitochondrial ATP production. Similar effects were recently found in populations of lake sturgeon, *Acipenser fulvescens*, exposed to increased water temperature during early development, a critical life stage in this species (Bugg, Thorstensen, Marshall, Anderson, & Jeffries, 2023). In our study, translation and cellular respiration suppression were accompanied by upregulation of several genes indicating heat-induced endoplasmic reticulum (ER) stress due to accumulation of unfolded or misfolded proteins in the ER. Genes involved in the calnexin/calreticulin (Cnx/Crt) cycle for protein folding quality control, together with key components of the ER-associated degradation pathway, were upregulated in offspring when reproduction occurred at elevated temperature. This indicates increased amounts of misfolded proteins in the ER, similar to findings in rainbow trout *Oncorhynchus mykiss* kidney following heat stress (Huang, Li, Liu, Kang, & Wang, 2018). Additionally, components of the unfolded protein response (UPR) pathway, activated to re-establish homeostasis when the ER folding capacity is overwhelmed, were upregulated in offspring because of reproduction at elevated temperature. One of these genes, for example, is the eukaryotic translation initiation factor 2α kinase 3 (EIF2AK3), a key stress sensor able to activate UPR and inhibit ribosome assembly and protein synthesis, leading to translation inhibition (Wek, Jiang, & Anthony, 2006). Upregulation of UPR markers and liver tissue damage was also found in largemouth bass *Micropterus salmoides* acutely exposed to elevated temperature (Zhao et al., 2022), while in mice livers ER stress markers and Cnx/Crt genes upregulation persisted for 21 days after a thermal injury, indicating long-term functional alterations of hepatic functions (Song, Finnerty, Herndon, Boehning, & Jeschke, 2009). Such prolonged suppression of protein synthesis and ATP production will likely result in energy limitations, fitness decrease and suboptimal growth (Sokolova, Frederich, Bagwe, Lannig, & Sukhotin, 2012). Indeed, offspring from parents that reproduced at elevated temperature were smaller at hatching and in worse body condition at 80 dph compared to offspring from control parents (Spinks et al., 2022). Therefore, reduced body weight of offspring from parents exposed to elevated temperature during reproduction could be linked to metabolic dysfunctions, suggesting either negative parental carry-over condition effects or detrimental effects of warming on embryogenesis. Exposure to elevated temperature during gametogenesis and embryo development, for example during a heatwave, seems therefore to cause long-term hepatic ER stress in the spiny damselfish, regardless of the offspring’s own post-hatching thermal environment, with lasting impairment of the translational and respiratory machineries.

We found that a mismatch between the thermal experience ontogenetic timings of parents deeply affected offspring gene expression profiles. Similarly, mismatches between parental developmental and reproductive temperatures differently affected routine oxygen consumption and body condition in third generation subadult *A. polyacanthus*, with detrimental effects (Bernal et al., 2021). Here, mismatching thermal environments across the parental lifetime, between development and reproduction, caused differential expression of many genes including the upregulation of genes involved in ER stress, Cnx/Crt cycle and UPR pathway. However, as such signals were absent when elevated temperature occurred during both parental development and reproduction, some of the detrimental effects of exposure to elevated temperature seem to be lessened. This is in line with the hypothesis that temporal autocorrelation between perceived environmental stimuli during successive sensitive ontogenetic windows might work as a positive feedback to reinforce the level of predictability of the future environment, ultimately affecting the reliability of the transmitted information and the adaptive nature of TGP (Bell & Hellmann, 2019; Burgess & Marshall, 2014; Leimar & McNamara, 2015). Despite some traits showing thermal acclimation when parental development and reproduction matched, exposure to elevated temperature throughout the parent lifespan seemed nevertheless to overall have detrimental effects on the subsequent generation. These offspring still showed suppression of translation and mitochondrial respiration, while also overexpressing genes involved in protein ubiquitination, indicating increased protein degradation, and apoptosis. Accordingly, reproduction did not occur when both parents experienced warming throughout their lives. Our results therefore indicate that *A. polyacanthus*, and perhaps other coral reef fishes, will struggle with the predicted increase in water temperature, with potentially serious adverse effects on population viability and persistence.

Persistence of organisms in a warming world will depend on their ability to acclimate, within and across generations, to the changing environment. In this study, we tackled the fundamental question of the importance of ontogenetic timing in the transmission of parental effects. We demonstrated that while a parental developmental experience of warming in *A. polyacanthus* might contribute to adaptive, although not anticipatory, transgenerational plasticity in their juvenile offspring, reproduction in warmer water will not have the same beneficial effects on the subsequent generation. Rather, if a heatwave should occur during the reproductive season, offspring may suffer from detrimental metabolic effects. Moreover, our results also suggest that parental pre-exposure to warming during development alters and potentially worsens the reproductive thermal signature. Finally, for the first time we show molecular evidence of physiological stress in offspring of parents with mismatching thermal experiences, suggesting that similar parental thermal histories are important for acclimation potential of their offspring. Overall, our results unveil new molecular mechanisms involved in transgenerational response to warming and demonstrate the importance of exposure timing of the previous generation’s environmental experiences to the capacity of individuals to cope with warmer ocean temperatures.

## Data accessibility

RNA-seq data for all individuals can be found under the BioProject PRJNA998209. All other data are provided in the electronic supplementary material.

## Authors’ contributions

L.C.B., R.K.S. and J.M.D. designed the experiment and collected the samples. L.C.B. and R.K.S. managed the fish rearing. L.C.B. prepared the samples for sequencing, analysed the sequencing data and wrote the first draft of the manuscript, with input from C.S.. J.M.D., P.L.M. and T.R. secured the funding. All authors read, provided comments and gave final approval for publication.

## Acknowledgements

We would like to thank the KAUST Bioscience Core Lab and the Marine and Aquaculture Research Facilities Unit at JCU for their help and support.

