## Supplementary Figures for "Timing specific parental effects of ocean warming in a coral reef fish"

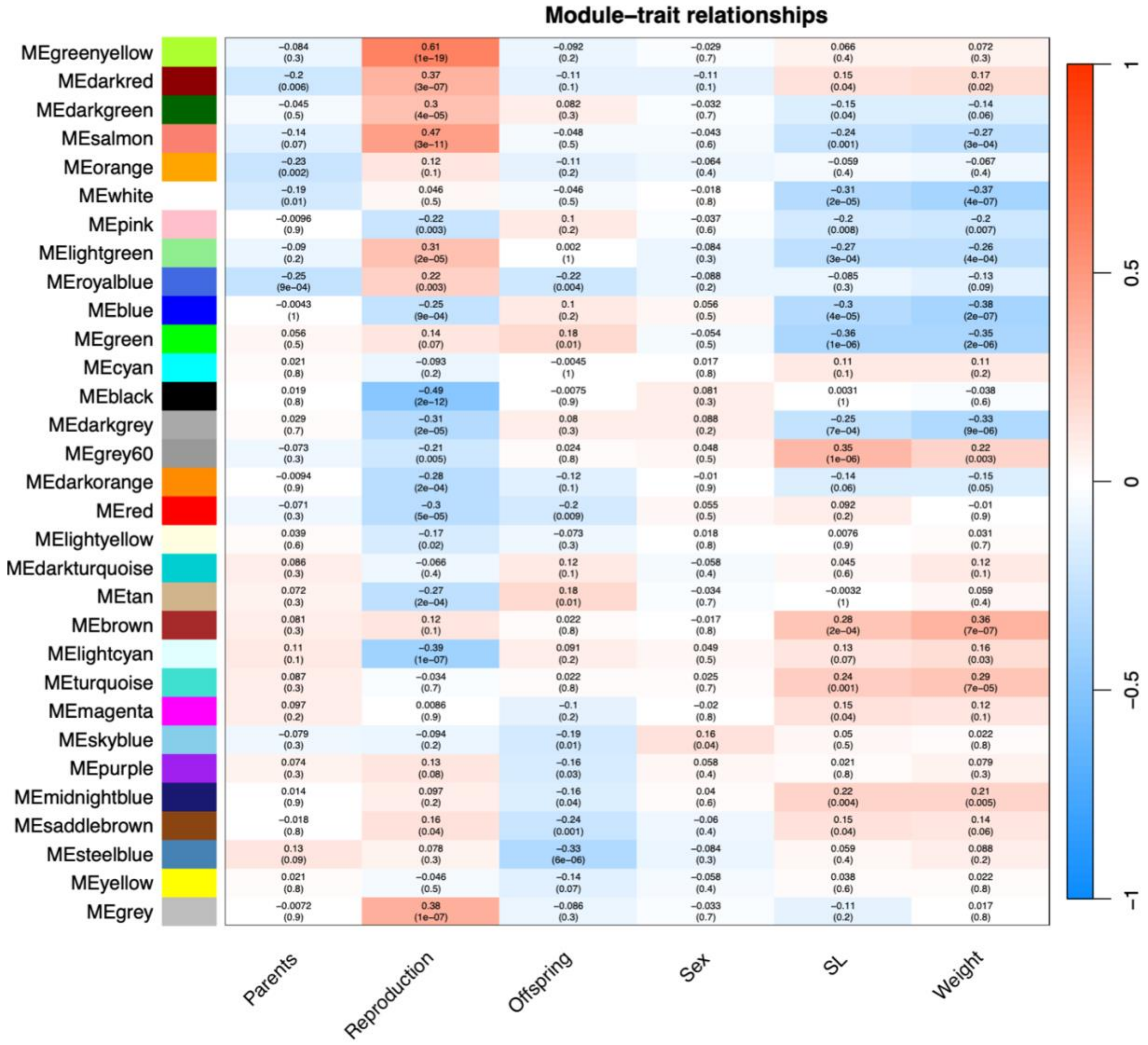


**Supplementary Figure S1. Gene cluster correlation with warming timing and offspring traits.** Heatmap of module-trait correlation analysis between detected gene network modules (denoted by an arbitrary color name) and fish treatments (elevated temperature during parental development - "Parents" , reproduction - "Reproduction", or offspring development - "Offspring") and F2 traits (sex, standard length - SL - and weight). In each cell is reported the correlation value for the corresponding module eigengenes and each treatment/trait and the p-value (in brackets).

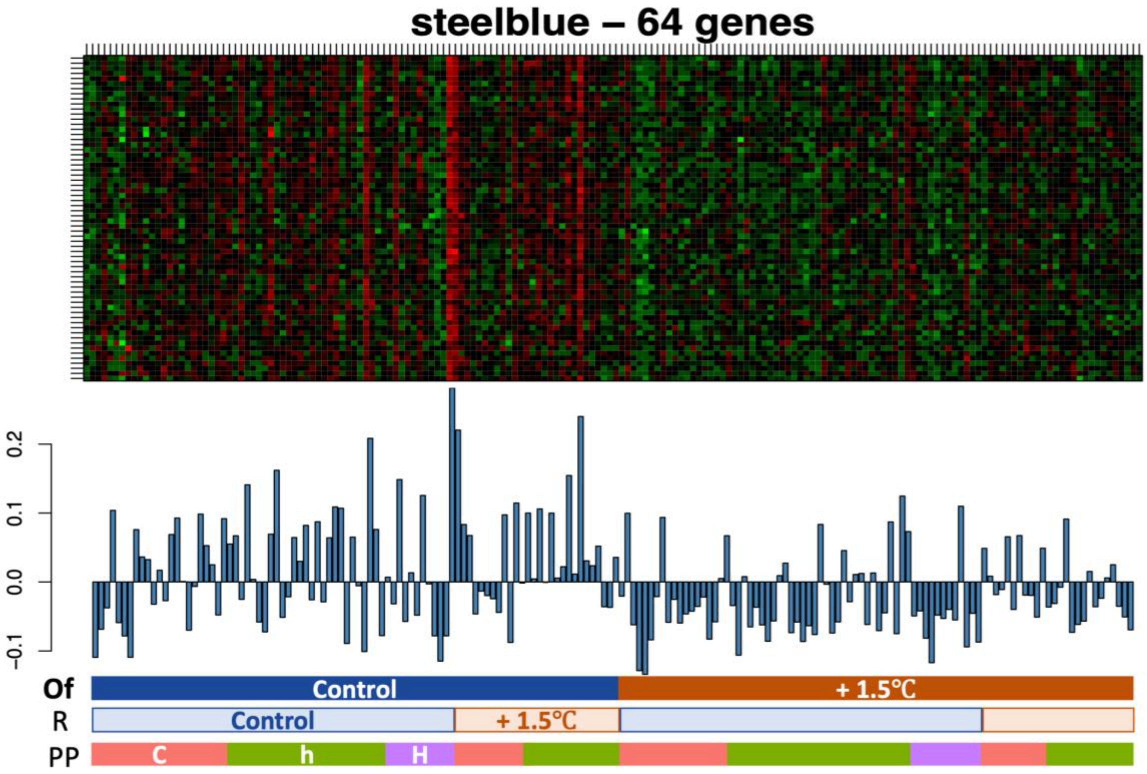


**Supplementary Figure S2. Heatmap of module correlated with offspring developmental temperature.** Eigenvalue of expression data for each sample in the steelblue module, negatively correlated with the offspring developmental thermal environment. Samples are arranged in the following order: by offspring developmental (Of), followed by reproductive (R) and then by parental pair developmental (PP) thermal treatments. In the parental pair development, "C" = both parents developed at control temperature, "h" = one parent developed at control temperature and one at +1.5°C, "H" = both parents developed at +1.5°C.

**PCAs with VST data**

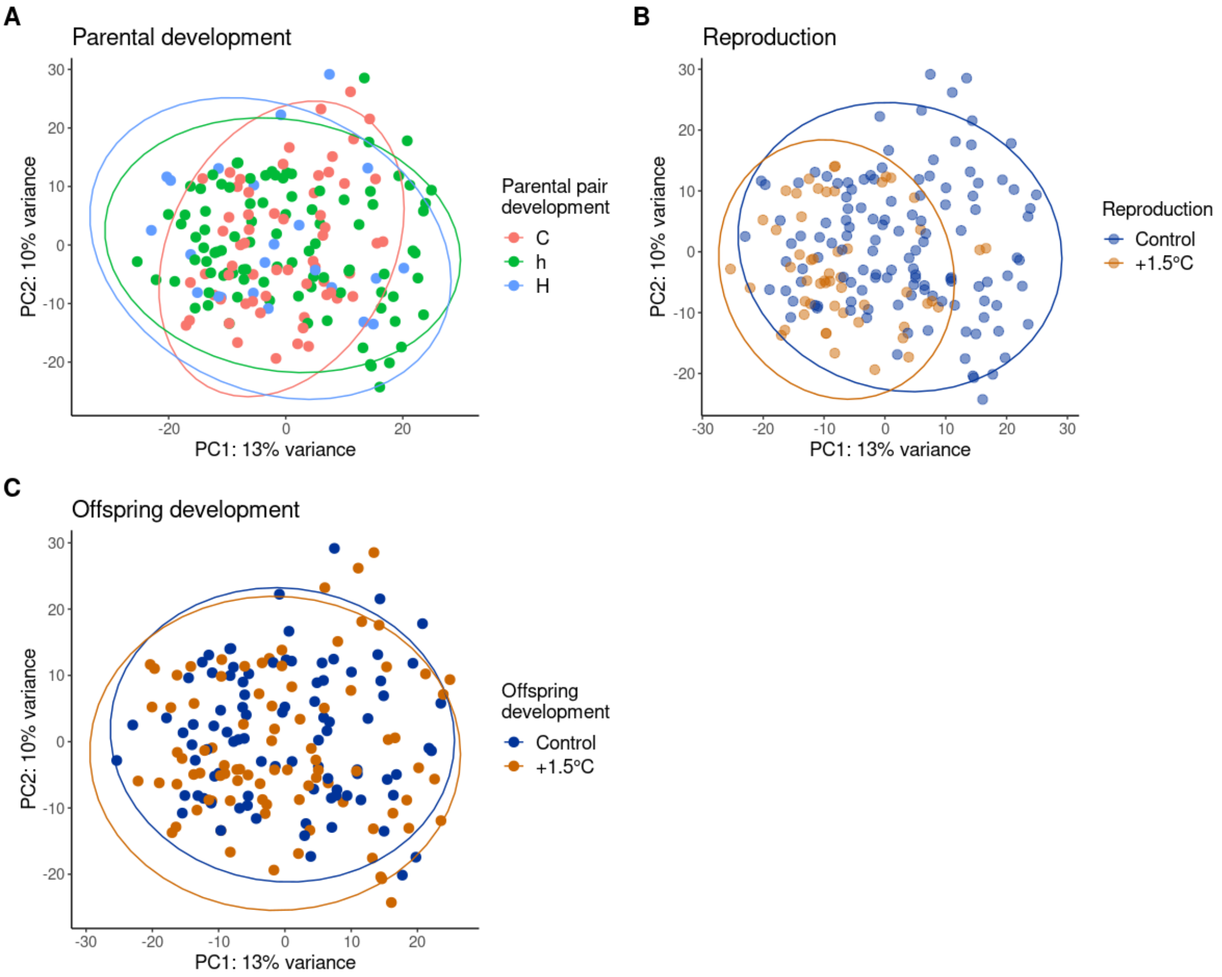


**Supplementary Figure S3. Principal component analyses (PCAs) of variance stabilized expression values of the 500 most variable genes.** Colors and ellipses of each plot represent different A) parental developmental B) parental reproductive and C) offspring developmental thermal environments. In the parental pair development, "C" = both parents developed at control temperature, "h" = one parent developed at control temperature and one at +1.5°C, and "H" = both parents developed at +1.5°C.

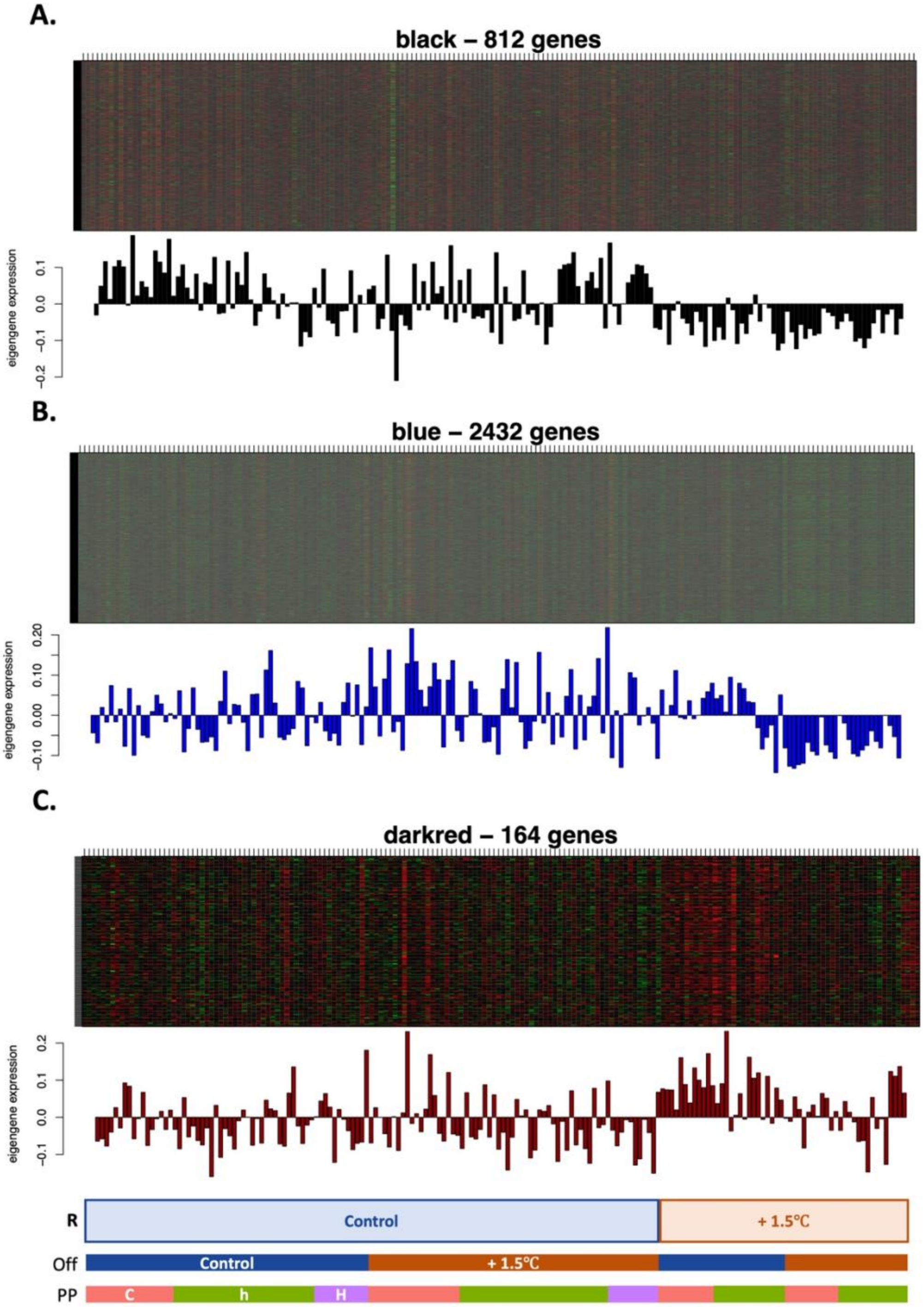


**Supplementary Figure S4. Heatmaps of modules correlated with reproductive temperature.** Eigenvalue of expression data for each sample in the A) black B) blue C) darkred modules, correlated with the reproductive thermal treatment. Samples are arranged in the following order: by reproductive (R), followed by offspring developmental (Off) and then by parental pair developmental (PP) thermal treatments. In the parental pair development, "C" = both parents developed at control, "h" = one parent developed at control and one at +1.5°C.
